# TRuML: A Translator for Rule-Based Modeling Languages

**DOI:** 10.1101/171306

**Authors:** Ryan Suderman, William S. Hlavacek

## Abstract

Rule-based modeling languages, such as the Kappa and BioNetGen languages (BNGL), are powerful frameworks for modeling the dynamics of complex biochemical reaction networks. Each language is distributed with a distinct software suite and modelers may wish to take advantage of both toolsets. This paper introduces a practical application called TRuML that translates models written in either Kappa or BNGL into the other language. While similar in many respects, key differences between the two languages makes translation sufficiently complex that automation becomes a useful tool. TRuML accommodates the languages’ complexities and produces a semantically equivalent model in the alternate language of the input model when possible and an approximate model in certain other cases. Here, we discuss a number of these complexities and provide examples of equivalent models in both Kappa and BNGL.

**CCS CONCEPTS:** • **Applied computing** → **Systems biology**; • **Computing methodologies** → *Simulation languages*;

## 1 INTRODUCTION

Rule-based modeling is a recently developed framework for modeling dynamics of biochemical reaction networks [2]. Its strength lies in the ability to encode large numbers of (or infinite) reactions with individual rules. In essence, parts of molecules that do not participate in a particular reaction are omitted from the rule, a paradigm known in the rule-based modeling community as *don’t care*, *don’t write.* This is analogous to the representation of reactions in organic chemistry in which a reaction involving some functional group may occur regardless of the configuration of the functional groups neighboring structure (*e.g.* R-OH represents the set of all alcohols). Rule-based frameworks can thus be used to build relatively concise models that exhibit considerable combinatorial complexity [12]. In simple cases (where the number of distinct biochemical species can be enumerated), rule-based models can be converted to a reaction network or system of equations and simulated using traditional methods (*e.g*. the stochastic simulation algorithm (SSA) [7] or numerical integration techniques). For cases involving systems that can generate large numbers of (or infinite) species, alternative kinetic Monte Carlo approaches based on the SSA have been developed that directly apply rules to a mixture of objects representing specific molecular configurations [3, 11]. In these approaches, rules serve as event generators where their iterative application to some initial mixture generates a stochastic trajectory.

## 2 RULE-BASED MODELING LANGUAGES

Underlying rule-based modeling frameworks are formal languages designed to encode biochemical interactions. Here, we are concerned with converting between two prominent rule-based modeling languages, the Kappa language [4] and the BioNetGen language (BNGL) [6]. These languages encode types of molecules and their interactions as human- and machine-readable plain text as a name and a list of *sites* that can interact with other molecules’ sites or occupy one of a finite number of predefined states (see the grammars in Appendix A). For example, given a molecule, or *agent*, ‘A’ with a site ‘b’ that only engages in binding other sites and a site ‘s’ that can occupy three distinct states (0, 1 or 2) or engage in binding, the BNGL encoding of the molecule type is

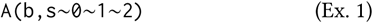

Writing rules involves constructing *patterns* composed of molecule types in which a subset of the molecule types’ sites are present in some specified state (including binding state). To clarify the terminology used here: patterns refer to language constructs that are used to match some set of biochemical species and *complexes* or *molecules* refer to the biochemical species themselves or to the specific objects that make up a pattern. The left-hand side (LHS) elements of a rule (denoted by patterns to the left of an arrow operator) identify the *reactants*, the right-hand side (RHS) elements identify the *products*, and the difference between the two sides is the representation of some physical or chemical transformation. Rules can then be applied during the course of simulation when the rule’s LHS matches (is *embedded* into) some biochemical species in the simulation mixture, precluding the need for *a priori* enumeration of the biochemical species that may arise. Consider a molecule ‘C’ with a single binding site ‘b’. A binding rule between ‘A’ from Ex. 1 and ‘C’ written in BNGL could then be

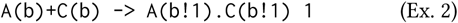

where ‘!1’ denotes the bond (‘1’ is the bond label, local to the rule), the ‘+’ operator denotes molecules that are not yet bound, the ‘.’ operator denotes molecules that are a part of the same complex (regardless of whether an explicit bond is present), and the trailing ‘1’ is the rule’s rate constant. This rule applies to any molecule ‘A’ that is not bound on site ‘b’ regardless of the state of ‘s’ or whether or not ‘s’ is bound to another molecule. Assuming other rules may be present that govern interactions for the site ‘s’ on ‘A’, some complexes that this rule could be applied to (*i.e.* that the ‘A’ molecule type in the rule’s LHS matches) are

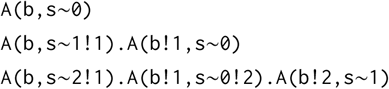

## 3 TRANSLATION

Here, we introduce TRuML, a **T**ranslator for **Ru**le-based **M**odeling **L**anguages that translates BNGL models into Kappa models and vice versa. Each framework contains a number of analytical tools and features that the other does not^1^ and modelers may wish to use both sets. The conversion can often be a nontrivial or error-prone exercise that is better left to automation. It is relevant to note that most of the difficult components to translate are semantic in nature, as the two languages are very similar lexically and syntactically. For clarity, our examples involve what we term *simple* rules and patterns that only involve agents or molecules and not other language features such as Kappa’s *tokens* or BNGL’s rule modffiers. To place our work in a broader context, work by Děd, *et al.* describes a third framework related to both Kappa and BNGL but operating at a higher level of abstraction to achieve the relation [5]. Additionally, the PySB framework for systems biology modeling in Python acts as a wrapper for features of both languages, facilitating use of rule-based modeling with other Python libraries for data analysis and visualization [10]. Outside of these examples, we are unaware of any other work that considers integration of or translation between these two rule-based modeling languages.

As a simple first example, we refer to Ex. 2 involving molecule ‘A’ and its interaction with molecule ‘C’. BNGL and Kappa have nearly equivalent syntax for defining a molecule’s name, its sites, and the states they may occupy (Grammars 1 & 3). However, one important distinction lies in the operators two languages use to connect multiple molecule. The Kappa equivalent to Ex. 2 is (under certain assumptions discussed in Section 3.3)

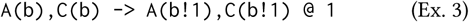

where the ‘@’ symbol separates the rule’s mechanism from its associated rate constant(s), distinct from BNGL’s whitespace separation. Note that, in Kappa, the ‘,’ operator connects all molecules types on one side of the rule, regardless of whether or not they are bound to each other. This difference from the ‘.’ and ‘ + ‘ operators has connotations for model translation that will be discussed in Section 3.3.

More subtle translation issues involve language conventions that are not obvious simply by reading a model. One example involves under-the-hood rate modification in BNGL-compatible simulation engines in order to maintain consistency with the law of mass action. If symmetries exist in a rule’s LHS then the rule’s rate is divided by the number of symmetries to account for the resulting combinatorial effect. Another way of describing this (as patterns in rule-based modeling languages can be represented as graphs) is to say that the rate is divided by the number of members in the automorphism group of the BNGL rule’s LHS. In contrast, automorphisms in Kappa rules must be explicitly accounted for. For example, given the BNGL rule:

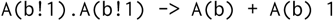

the equivalent Kappa rule would be

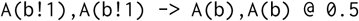

since there are two automorphisms in the rule’s LHS.

### 3.1 Semantic objects

TRuML parses rule-based models written in either language and converts the plain-text representations into semantic classes that generate appropriate text in either BNGL or the Kappa language. For example, both Kappa and BNGL require molecule type definitions as in Ex. 1. A number of other features of the languages are directly analogous (see Appendix A.1 & Table 1). Some examples include:

- Initial conditions that specify numbers of biochemical species at the start of a simulation
- Observables^2^ that track the number of a particular pattern throughout the simulation
- Expressions that represent both static^3^ and dynamic^4^ quantities. Kappa uses *variables* to define both, while BNGL distinguishes between static quantities (*parameters*) and dynamic quantities (*functions*)

**Table 1:**
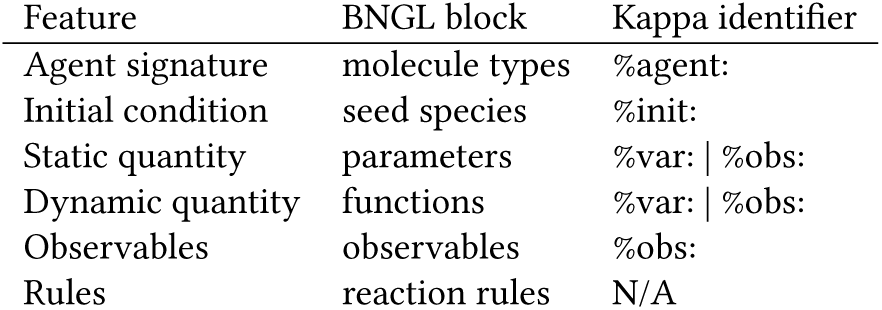
Defining analogous features in BNGL and Kappa. The ‘|’ symbol denotes that either Kappa identifier can be used.

### 3.2 Identical site names

One major difference between the two languages is the ability to define molecule types with identical sites in BNGL, which is not allowed in Kappa. In certain biological systems (*e.g*. mast cell signaling networks) the relevant molecules may exhibit multiple independent and functionally identical structures (*e.g*. antibodies with two antigen-binding *Fab* arms), and the BioNetGen software suite was designed to accommodate these cases by allowing identically named sites and appropriately scaling the rate constants using statistical factors. TRuML converts the BNGL molecule representation with identical sites to a similar Kappa representation with numbered sites that behave identically. For example, a bivalent antibody (‘Ab’) molecule and a trivalent antigen (‘Ag’) might be encoded in BNGL as follows:

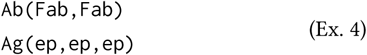

where the ‘Fab’ sites on an antibody bind the epitopes (‘ep’ sites) on an antigen molecule. TRuML renames the sites systematically for translation to Kappa

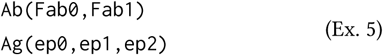

Rules must also be appropriately modified. A binding rule between free antigen and antibody (as defined in Ex. 4) in BNGL would be^5^:

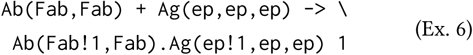

whereas the equivalent statement in Kappa must consider all 6 site permutations from the translated molecule definitions (in Ex. 5) resulting:

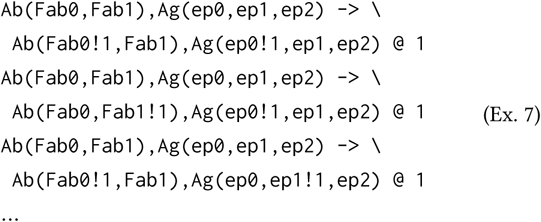

Note that the BNGL rule’s rate will be scaled by a statistical factor by the simulation engine as there are 6 different interactions governed by this rule. Not coincidentally, our convention for translation to Kappa involves 6 rules that are applied using the unscaled rate.

Also affected are patterns that define observables in terms of identically named sites. These observables must have their Kappa equivalents multiplied by the number of automorphisms in the BNGL pattern, since the Kappa pattern will only have the trivial automorphism. While BNGL automatically adjusts rates to account for the automorphism group of a rule’s LHS, the BNGL observable counts are not similarly scaled. As might be expected, translation becomes increasingly complex if identically named sites in one molecule can bind identically named sites in another such as the following rule where an antigen ‘crosslinks’ multiple antibodies, meaning it is already bound to at least one antibody when binding another antibody:

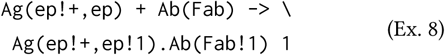

Here the ‘!+’ notation means that a bond on this site is required for a pattern match, but the binding partner is irrelevant.

### 3.3 Molecularity

Perhaps most complex issue to handle in converting between Kappa and BNGL is molecularity (*e.g.* determining whether or not molecules are bound). Rules involving molecules in the same or distinct complexes are easily identified in BNGL syntax due to the dual ‘.’ and ‘+’ operators.^6^ In Kappa this is a bit more difficult, as the absence of an explicit bond between two molecules does not preclude a pattern from matching two molecules that are a part of the same complex. For example, the Kappa pattern A(b),A(b) embeds into the complex A(b,s~θ!1),A(b,s~θ!1).

This is termed *ambiguous molecularity* and modelers using the Kappa language are advised to avoid this if possible by including additional context into rules where this might occur. The rule in question could be rewritten as two rules to clarify the modelers’ intentions:

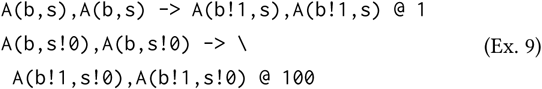

If ambiguity is unavoidable, as in cases involving polymerization where rings must not form (Ex. 8), Kappa modelers may use the rate syntax “k1 {k2}” (see Grammar 2) at the expense of simulation performance as in the following rule:

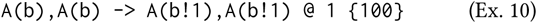

The first number (k1) is the reaction rate when the molecule types on the rule’s LHS match separate molecules (intermolecular association; the first rule in Ex. 9) and the second number in brackets (k2) is applied when the matched molecules are in the same complex (intramolecular association; the second rule in Ex. 9). TRuML detects this syntax and will appropriately construct two forms of the above rule in the BNGL syntax:

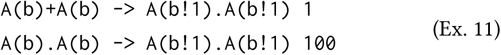

Note that the second rule’s LHS contains an implicit bond as the two ‘A’ molecule types are connected with the ‘.’ operator. When converting from BNGL to Kappa, it is not always obvious if intramolecular bonds can form. If TRuML cannot make this determination automatically, it will include two separate rules for each Kappa binding rule, one with each operator. The user may also specify the absence of any intramolecular bond formation when converting from Kappa to BNGL, and only the ‘+’ operator will be used for binding rules.

Translating rules from BNGL to Kappa can be considerably more difficult, due to the nature of the BNGL ‘+’ and ‘.’ operators. Each rule translated into Kappa must guarantee that the molecularity constraints placed on the rule by the BNGL operators hold for the Kappa rule. As a result, some BNGL rules cannot be converted to Kappa rules. For example, BNGL models containing rules with greater than 2 ‘+’ operators on either the LHS or RHS of the rule cannot be translated into Kappa. Furthermore, some rules such as the following fall outside the purview of Kappa’s ambiguous molecularity rate notation and must be clarified by adding explicit binding context if possible:

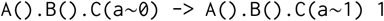

This notation cannot be translated when the implicitly bound pattern can match polymers of arbitrary size. TRuML therefore has a user-specified flag that tells the application if polymers or rings can form. This information will prevent potential infinite loops in the pattern disambiguation process.

## 4 RESULTS

In this section, we compare simulation results of translated models to the original models. These translations are equivalent in the sense that no information is lost in the encoding of the reaction network (no approximations are present in the translation). Both sets of results use Kappa’s KaSim engine [1] and BNGL’s NFsim engine [11] for simulation. Since both use stochastic simulation methods, the results shown from either language will not be identical, but will be statistically indistinguishable.

### 4.1 Futile cycle model

As a simple demonstration of TRuML, we developed a rule-based model of a futile cycle written in the Kappa language. It is composed of 3 molecule types: two enzymes (E1 & E2) and a substrate (S). One enzyme modifies the substrate and the other removes the modification, both according to the Michaelis-Menten model of enzyme kinetics. In Figure 1 we plot timecourse data from the original and translated models, revealing statistically equivalent simulation trajectories.

**Figure 1:**
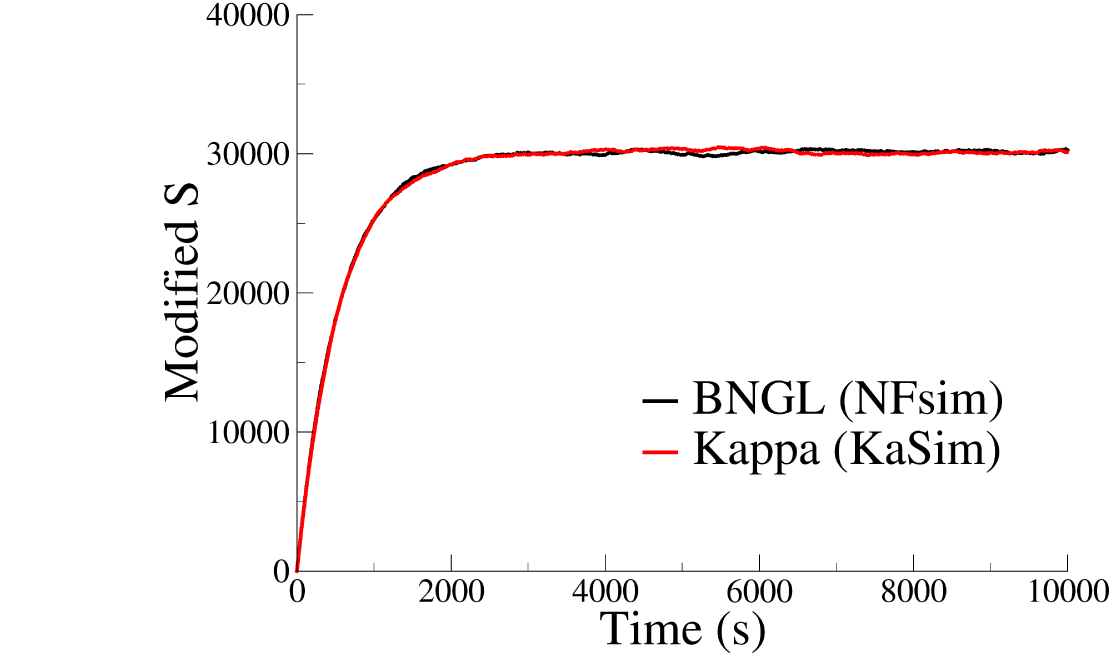
Dynamics for the number of modified substrate observables regardless of binding state in a Kappa futile cycle model (red) and its BNGL translation (black).

### 4.2 Antibody-antigen binding model

As a second example, we constructed a BNGL model of trivalent antigen molecules binding bivalent antibody molecules based on Ex. 4, Ex. 6, and Ex. 8. The BNGL model contains three rules and two molecule types, but because both molecule types have identical sites and the rules are capable of generating polymers, the resulting Kappa translation has a notably larger set of rules. To further demonstrate that equivalent models can be represented using different conventions, we then translated the Kappa model back into BNGL. The dynamics for fully-bound antigen from all three models are shown in Figure 2.

**Figure 2:**
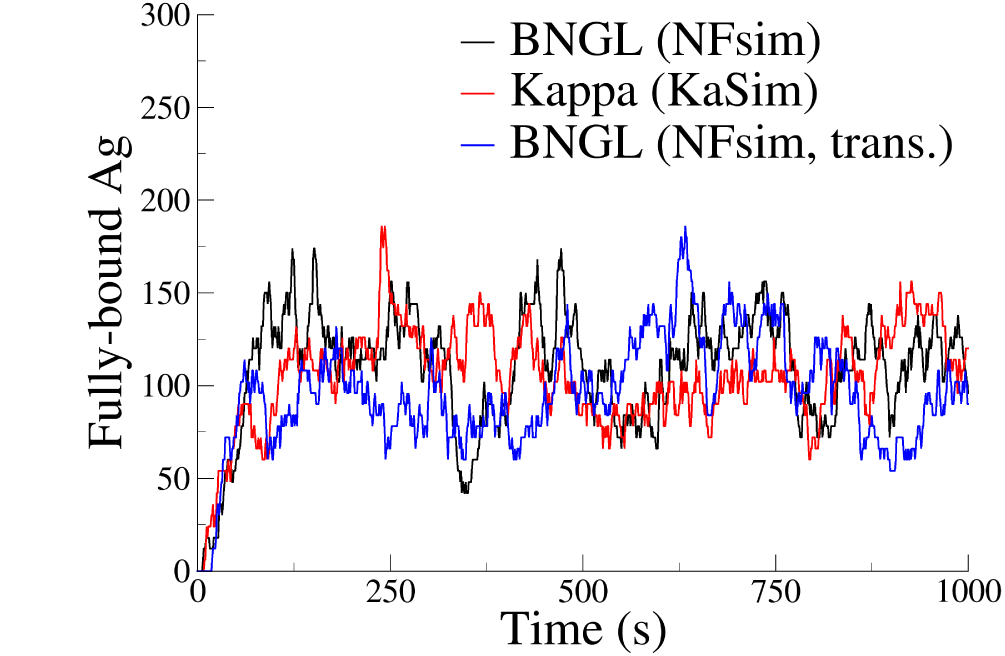
Dynamics for the fully-bound antigen observable “Ag(ep! +, ep! +, ep!+)” in a BNGL antigen-antibody model (black), its Kappa translation (red) and a second translation back into BNGL (blue). Note that this is the number of *patterns* (6x the number of actual molecules corresponding to this pattern).

We also recorded the system state for each simulation after 1000 seconds of simulation time to analyze the distribution of aggregates formed. Each simulation generated between 125 and 135 unique molecular species (not counting the remaining antigen and antibody monomers) and between 350 and 380 total complexes, from a starting point of 6000 monomeric antigen molecules and 600 monomeric antibody molecules. We generated histograms for each trajectory (seen in Figure 3) and performed pairwise Kolmogorov-Smirnov tests on the distributions. All of the tests failed to distinguish between the distributions.

**Figure 3:**
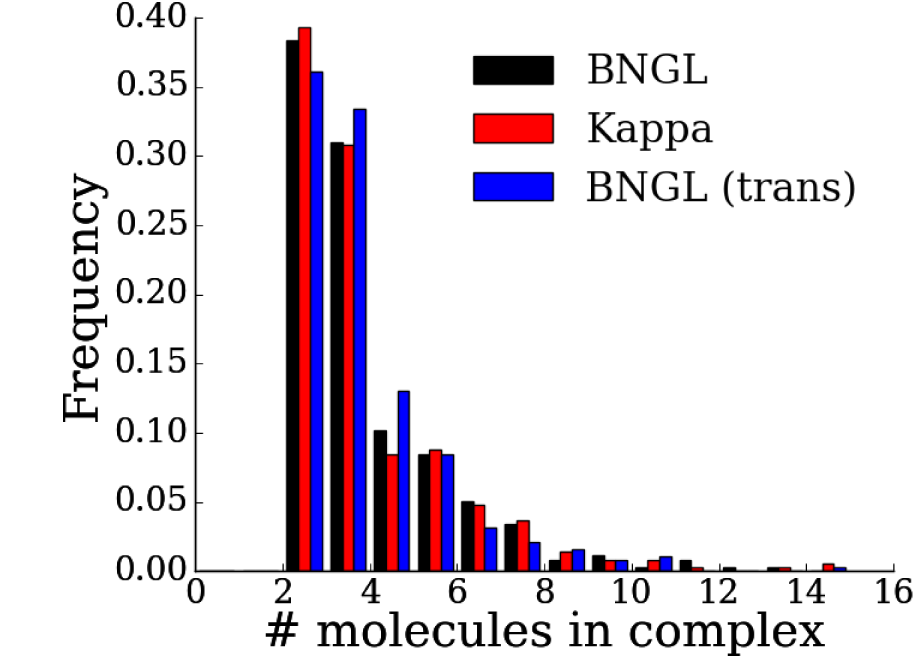
A histogram showing the frequency of aggregate size (omitting monomers) at the end of a 1000-second simulation for each antigen-antibody model. Colors match those in Figure 2

## 5 DISCUSSION

As with translating natural language, it is important to note that a statement in one language may have numerous equivalent statements in another. As a result, translating from one language to the other and back again may produce a model that appears distinct from the original despite its semantic equivalence. Furthermore, TRuML utilizes one particular (arbitrarily chosen) convention for translating between these rule-based modeling languages. An alternative approach to representing molecules with identically named sites in Kappa has been suggested, which involves a complex that does not dissociate [8]. Using this convention, the antigen molecule in Ex. 4 could be represented with the following molecule type and complex:

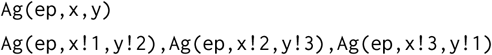

In addition to representing three identically named sites, this structure also preserves the implicit symmetry in its BNGL equivalent through its symmetric bonding structure. Certain rules could then be written in a more BNGL-like style, especially if the antibody molecule type follows a similar convention.

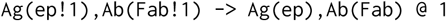

Finally, both languages are capable of describing models that are impossible to exactly translate into the other language. Some of the issues involving rule structure resulting in incompatible models have already been discussed (*e.g*. implicit bonds that match polymers). However other features of both frameworks preclude exact translation as well. BNGL models can contain a specific type of observable called a *Species* observable, which tracks the number of complexes in the mixture that contain a particular pattern. This is typically used for determining the number of aggregates on-the-fly. On the other hand, Kappa models allow variables to take an infinite value. Encoding a rule that takes place instantly (in simulation time) is easily done by assigning it an infinite rate. Only an approximation of this can be reached in BNGL (typically by assignment of a very large rate constant).

As the readers may have noticed, the complexities in translating between BNGL and Kappa generally come when converting BNGL models with certain properties to Kappa models. Features that allow concise representation of certain types of systems in BNGL can occasionally result in Kappa models with considerably more rules upon translation. This is not to say that BNGL is more useful than Kappa (or vice versa) when considering the entire modeling framework associated with these languages. Indeed, Kappa is arguably more tractable for formal analysis as outlined in [8]. The occasionally non-trivial differences between these two languages make TRuML useful for systems biology modelers who wish to fully exploit the features of both the Kappa and BNGL frameworks.

## A GRAMMARS AND OTHER SYNTAX

### A.1 Definition Syntax

Syntax for defining equivalent features differs slightly between the two formalisms. Consider how the two languages represent mathematical expressions involving time-varying quantities:

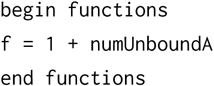

where *numUnboundA* is defined in an observable block. The equivalent Kappa definition is simply the line:

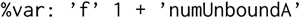

where *numUnboundA* is similarly defined as an observable. Table 1 defines equivalent features in BNGL and Kappa.

### A.2 Grammars

Grammars describing patterns and simple rules in Kappa and BNGL are given in Extended Backus-Naur Form. Character sequences in single quotes are terminal symbols, and the grammars use the following non-terminal symbols defined by regular expressions:

- 〈*integer*〉 = [0-9]+
- 〈*bName*〉 = [a-zA-Z][a-zA-Z_0-9]^∗^
- 〈*kName*〉 = [a-zA-Z][a-zA-Z_0-9+-]^∗^
- 〈*string*〉 = .^∗^
- 〈*newline*〉 = \n
- 〈*ws*〉 = [ \t]

The grammars do not include treatment of whitespace (except for that needed to separate the products from the rate in BNGL) or the line continuation character (\) for simplicity.

**Grammar 1:**
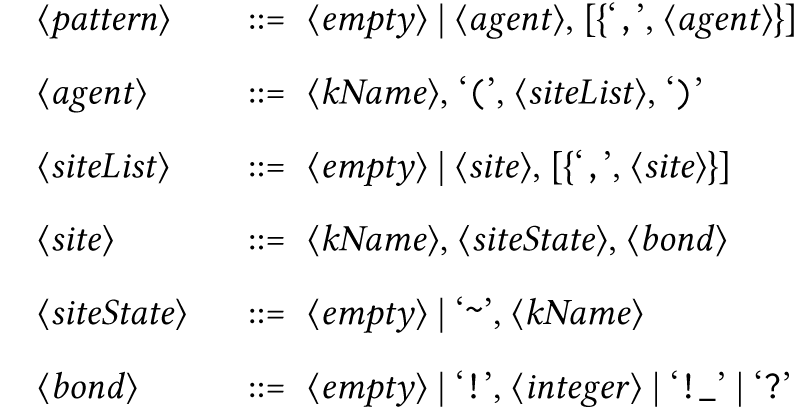
Kappa patterns as defined in [1]

**Grammar 2:**
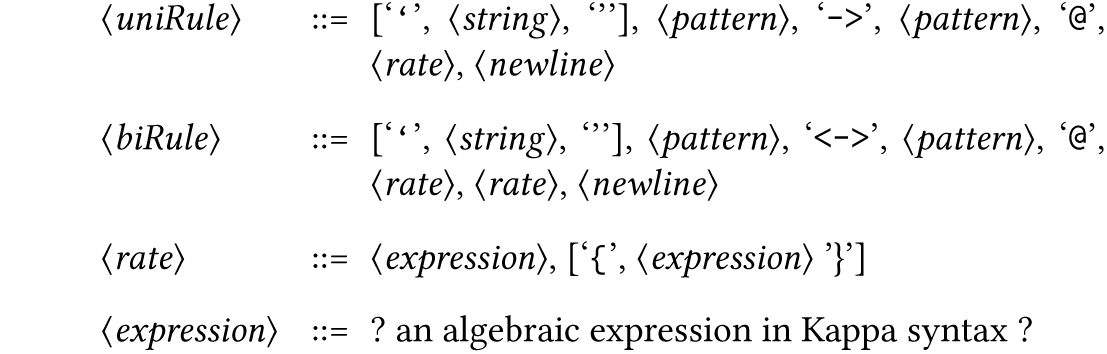
A grammar for simple rules in Kappa (using patterns from Grammar 1) based on grammars in [1]

**Grammar 3:**
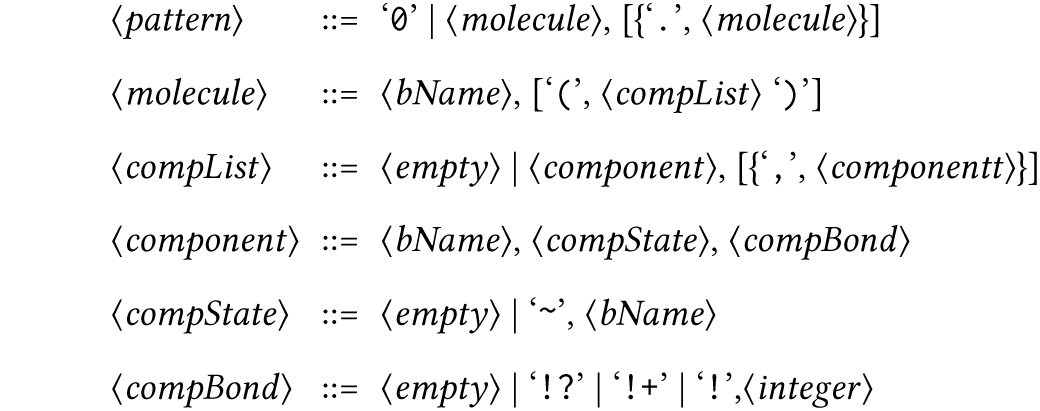
BNGL patterns, based on the grammar defined in [9]. ‘0’ denotes the empty pattern. Note that BNGL patterns only match molecules that are in the same complex.

**Grammar 4:**
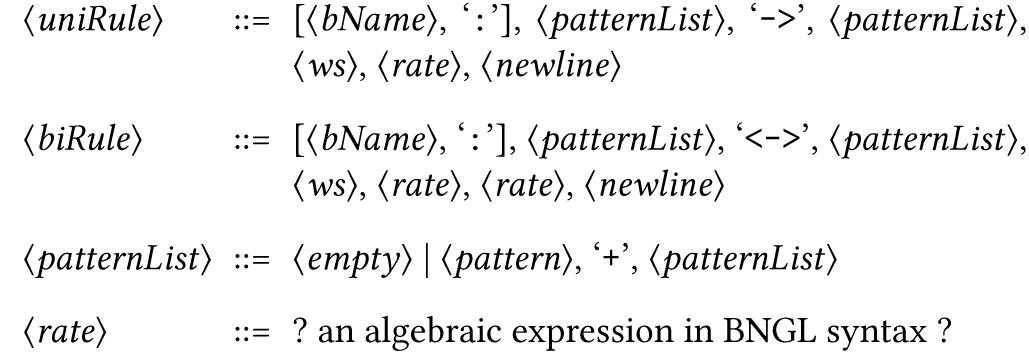
A grammar for simple rules in BNGL using patterns from Grammar 3 and based on the grammar in [9].

## B TOOLS FOR MODEL AND SIMULATION ANALYSIS

Both languages are accompanied by software that enables simulation and analysis of the model itself. Web resources can be found for both Kappa (http://dev.executableknowledge.org/) and BNGL (http://bionetgen.org/index.php/Main_Page)

### B.1 Kappa

The program KaSim simulates models written in the Kappa language. In addition to the language describing the interactions between biomolecules, KaSim also contains a *perturbation* language exists that enables the modeler to specify modifications to the simulation state on-the-fly. One feature of the perturbation language is the ability to flag an observable in order to record the sequence of events that leads to its formation. Kappa models can also be analyzed without simulation (termed *static* analysis) using the KaSa program. KaSa can be used to build the *contact map*, which describes how molecules in the model may be connected to each other. Another is the influence map, which describes how the execution of a rule during simulation may affect the propensity of other rules to be executed.

### B.2 BNGL

The BioNetGen software suite (or simply BioNetGen) has a number of methods for model simulation, and BNGL models typically contain an *actions* block with commands that engage simulation. Two methods require generating a *network file* (hence the name ‘BioNetGen’), a process that enumerates all species a particular rule set can form, assuming polymerization cannot occur. The network file is interpreted either as a system of differential equations or a Markov chain and is simulated accordingly. The previously mentioned NFsim (Network-Free simulation) engine is used for on-the-fly generation of molecular species and is distributed with accompanying scripts for analyzing results in MATLAB. BioNetGen also contains visualization tools that generate graph-based images of the set of rules or visual model summaries.

## ACKNOWLEDGMENTS

This work was supported by NIH/NIGMS grant R01 GM111510 and the Center for Nonlinear Studies at Los Alamos National Laboratory.

## Footnotes

1 See Appendix B

2 BNGL has a certain type of observable that is incompatible with Kappa (Section 5)

3 Examples of static quantities are interaction affinities or protein copy numbers

4 Dynamic quantities are typically calculations involving the number of embeddings of some pattern at each time step in the simulation, such as the Michaelis-Menten approximation of enzyme activity

5 The backslash is a line-continuation character

6 The NFsim simulation engine currently does not check for product molecularity while the built-in BioNetGen SSA and numerical integration algorithms do. TRuML includes a flag for conversion to either of these implementations

## REFERENCES

[1] Pierre Boutillier, Jérôme Feret, Jean Krivine, and Lý Kim Quyên. 2017. KaSim and KaSa Reference Manual. http://dev.executableknowledge.org/docs/KaSim-manual-master/KaSim_manual.htm.

[2] Lily A. Chylek, Leonard A. Harris, Chang-Shung Tung, James R. Faeder, Carlos F. Lopez, and William S. Hlavacek. 2014. Rule-based modeling: a computational approach for studying biomolecular site dynamics in cell signaling systems. Wiley Interdisciplinary Reviews. Systems Biology and Medicine 6, 1 (2014), 13–36. DOI: https://doi.org/10.1002/wsbm.1245 arXiv: NIHMS150003

[3] Vincent Danos, Jérôme Feret, Walter Fontana, and Jean Krivine. 2007. Scalable Simulation of Cellular Signaling Networks. In Programming Languages and Systems. Springer Berlin Heidelberg, Berlin, Heidelberg, 139–157. DOI: https://doi.org/10.1007/978-3-540-76637-7_10

[4] Vincent Danos and Cosimo Laneve. 2004. Formal molecular biology. Theoretical Computer Science 325, 1 (2004), 69–110. DOI: https://doi.org/10.1016Zj.tcs.2004.03.065

[5] T. Děd, D. šafránek, M. Troják, M. Klement, J. šalagovič, and L. Brim. 2016. Formal Biochemical Space with Semantics in Kappa and BNGL. Electronic Notes in Theoretical Computer Science 326 (2016), 27–49. DOI: https://doi.org/10.1016/j.entcs.2016.09.017

[6] James R. Faeder, Michael L. Blinov, and William S. Hlavacek. 2009. Rule-Based Modeling of Biochemical Systems with BioNetGen. In Systems Biology, Ivan V. Maly (Ed.). Humana Press, Totowa, NJ, 113–167. DOI: https://doi.org/10.1007/978-1-59745-525-1_5

[7] Daniel T Gillespie. 2007. Stochastic simulation of chemical kinetics. Annual Review of Physical Chemistry 58 (2007), 35–55. DOI: https://doi.org/10.1146/annurev.physchem.58.032806.104637

[8] Russ Harmer, Vincent Danos, Jérôme Feret, Jean Krivine, and Walter Fontana. 2010. Intrinsic information carriers in combinatorial dynamical systems. Chaos 20, 3 (2010), 1–16. DOI: https://doi.org/10.1063/1.3491100

[9] Justin S Hogg. 2013. Advances in Rule-based Modeling: Compartments, Energy, and Hybrid Simulation, with Application to Sepsis and Cell Signaling. (August 2013). http://d-scholarship.pitt.edu/19621/

[10] Carlos F Lopez, Jeremy L Muhlich, John A Bachman, and Peter K Sorger. 2013. Programming biological models in Python using PySB. Molecular Systems Biology 9, 646 (2013), 646. DOI: https://doi.org/10.1038/msb.2013.1

[11] Michael W Sneddon, James R Faeder, and Thierry Emonet. 2011. Efficient modeling, simulation and coarse-graining of biological complexity with NFsim. Nature Methods 8, 2 (2011), 177–183. DOI: https://doi.org/10.1038/nmeth.1546

[12] Ryan Suderman and Eric J. Deeds. 2013. Machines vs. Ensembles: Effective MAPK SignalingthroughHeterogeneousSets ofProtein Complexes. PLoSComputational Biology 9, 10 (2013).

